# Distinct duration and diet dependent lipid profiles and renal complications in GCK-MODY

**DOI:** 10.1101/2024.09.17.613598

**Authors:** Yadi Huang, Yuxin Fan, Yang Liu, Xuan Liu, Wei Li, Ying Yang, Ziyue Zhang, Shifeng Ma, Shuhui Ji, Shanshan Chen, Hua Shu, Wenli Feng, Kunling Wang, Qing He, Wenjun Qi, Yu Fan, Xin Li, Ming Liu

**Affiliations:** Department of Endocrinology and Metabolism, Tianjin Medical University General Hospital, Tianjin, 300052, China; NHC Key Laboratory of Hormones and Development, Chu Hsien-I Memorial Hospital and Institute of Endocrinology, Tianjin Medical University, Tianjin, 300134, China

## Abstract

Heterozygous inactivating mutations in the glucokinase (GCK) gene cause one of the most common types of maturity-onset diabetes of the young type 2 (MODY2), also named GCK-MODY. Studies suggest that, unlike other types of diabetes, patients with GCK-MODY do not have increased risk of diabetic complications and therefore do not typically require antihyperglycemic therapy. However, long-term outcomes of GCK-MODY on lipid profiles and chronic complications under different dietary scenarios remain unclear. Herein, using a knock-in mouse model expressing a novel MODY causing mutation GCK-Q26L (GCK^Mut^), we examined age- and diet-related lipid profiles and diabetic complications. We found that, although GCK^Mut^ mice exhibited mild elevated blood glucose, the lipid profiles, body fat composition, and diabetic kidney disease (DKD) were initially improved in high-fat-diet (HFD) fed mice at the age of 28 weeks, supporting potential beneficial effects of GCK^Mut^ on lipid metabolism and kidney health. Unexpectedly however, those protective effects diminished by 40 weeks, and became more severe dyslipidemia and kidney injury associated with renal fibrosis and inflammation at 60-week-old mice fed with normal diet (ND) or HFD. Those findings revealed distinct duration- and diet-dependent effects of inactivating GCK mutation on lipid profiles and DKD, highlighting previously unrealized long-term chronic complications in GCK-MODY.

## Introduction

Glucokinase (GCK), a glycolytic enzyme belonging to the hexokinase family, plays a pivotal role in catalyzing the conversion of glucose into glucose-6-phosphate, a primary step in glucose metabolism within pancreatic β-cells and hepatocytes (1–3). Heterozygous inactivating mutations in the GCK gene cause maturity-onset diabetes of the young type 2 (MODY2, also named GCK-MODY), which is one of the most common types of monogenic diabetes (4–6). Patients with GCK-MODY typically present mild to moderate non-progressive elevated fasting blood glucose (FBG) without significant abnormalities or even improvement of lipid metabolism (7, 8). More importantly, current evidence indicates that individuals with GCK-MODY are not associated with increased risks of chronic diabetic complications compared to the other types of diabetes (9). Therefore, current consensuses suggest that such patients may not necessitate pharmacological interventions (10, 11).

However, due to the limited number of patients diagnosed with GCK-MODY, there is a paucity of prospective interventional or observational studies examining the long-term outcomes of GCK-MODY under different dietary conditions. Additionally, given the potential impact of persistent hyperglycemia on ocular, vascular, neurological and renal diseases (12–15), it is important to investigate long-term effects of inactivating GCK mutations on lipid metabolism and functional and structural alterations in microvascular complications associated with prolonged hyperglycemia. Recently, we have established a knock-in mouse line expressing a novel GCK mutation (GCK-Q26L, GCK^Mut^) identified in the family with early onset diabetes, and demonstrated that this mutation indeed impaired insulin secretion causing mild elevated FBG and impaired glucose tolerance that mirrors the phenotypes of patients with GCK-MODY (https://doi.org/10.1101/2024.08.13.24311668). In this study, using this knock-in mouse model, we examined age- and diet-related lipid profiles and diabetic complications in the GCK-MODY. We found that GCK Mutant indeed presented beneficial effects on lipid metabolism, body fat composition, and renal injury in high-fat-diet (HFD) fed GCK^Mut^ mice at the age of 28 weeks. However, those protective effects diminished with time, and more severe dyslipidemia and kidney injury associated with renal fibrosis and inflammation were observed in GCK^Mut^ mice fed with HFD at the age of 60 weeks. Those findings revealed distinct duration- and diet-dependent effects of inactivating GCK mutation on body composition, lipid profiles, and diabetic kidney disease (DKD), highlighting previously unrealized long-term metabolic abnormalities and chronic diabetes complications in GCK-MODY. This is poised to offer valuable insights for the prevention and treatment of individuals diagnosed with GCK-MODY aiming to improve long-term reduction of chronic complications. Given the fact that decreased expression and function of GCK are also seen in type 2 diabetes (16), this study may also have clinical implications in the management of type 2 diabetes (T2D).

## Results

### GCK^Mut^ impairs insulin secretion, but improves lipid profiles in early normal diet (ND) mice

We have monitored body weight (BW) and FBG of ND fed mice every 4 weeks until 60 weeks of age, and found that GCK^Mut^ caused non-progressively mild elevated FBG without affecting BW (Fig. 1A-B). To evaluate the effect of GCK^Mut^ on insulin secretion and glucose tolerance, we performed intraperitoneal glucose tolerance tests (IPGTT) at the ages of 28, 40, and 60 weeks, respectively. We found that GCK^Mut^ resulted in impaired glucose tolerance associated (Fig. 1C-E, Fig. S1A-C) with decreased basal and glucose stimulated insulin secretion (Fig. 1F-H, Fig. S1D-F). However, GCK^Mut^ did not appeared to affect insulin sensitivity during intraperitoneal insulin tolerance tests (IPITT) (Fig. 1I-K, Fig. S1G-I). Next, we examined body composition and lipid profiles of GCK^Mut^ mice, and found that although GCK^Mut^ did not change the body fat composition, the weight of perirenal white adipose tissue (pWAT), and the weight of gonadal white adipose tissue (gWAT) (Fig. 1L-N), it indeed decreased levels of serum triglyceride (TG) and low-density protein cholesterol (LDL-c) in 28-week-old mice. However, those beneficial effects on lipid metabolism diminished at 40 and 60 weeks of age (Fig. 1O-R). Those data demonstrate that decreased activity of GCK causes mild non-progressive hyperglycemia with improved lipid profiles at young age and this improvement waned over time.

**Figure 1.**
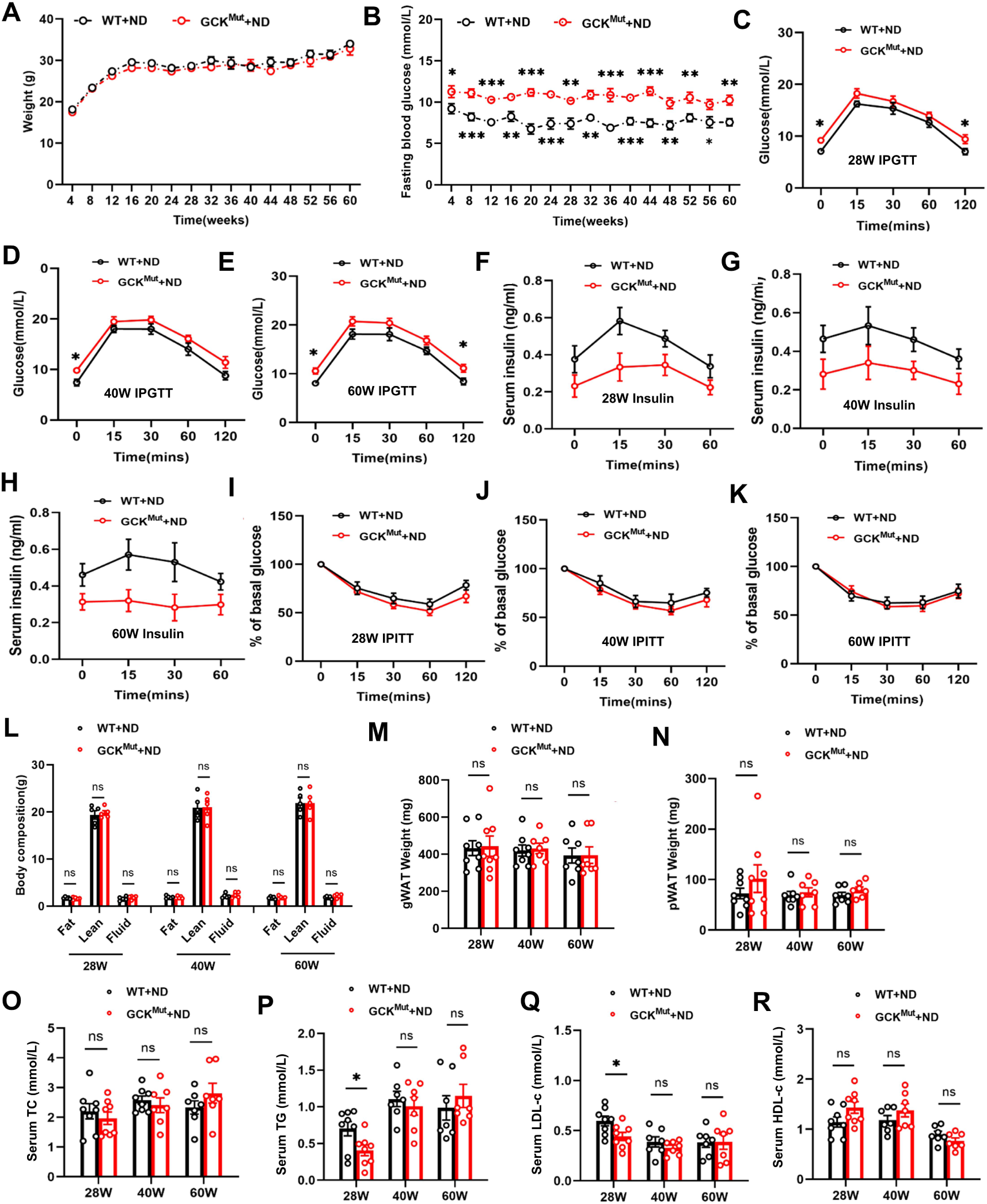
GCK^Mut^ mice had elevated FBG but improved lipids. Body weight (**A**) and fasting blood glucose (**B**) were monitored every 4 weeks (n=7-8 mice/group). (**C-E**) Two groups of mice were intraperitoneally injected with 1 g/kg for IPGTT at 28, 40, or 60 weeks of age (n=5mice/group). (**F-H**) Serum insulin levels were measured by ELISA during IPGTT in both groups of mice (n=5 mice/group). (**I-K**) Two groups of mice were subjected to IPITT (intraperitoneal injection of 0.75 U/kg, n=5mice/group) at 28, 40, or 60 weeks of age. (**L**) Body composition analysis of both groups of mice using a body fat analyzer. (**M-N**) Wet weights of fresh pWAT and gWAT of mice were measured at 28, 40, and 60 weeks of age. Blood samples were used to measure lipid levels including TC (**O**), TG (**P**), LDL-c (**Q**), and HDL-c (**R**), n=5-8mice/group. Values are expressed as mean±SEM. WT+ND vs. GCK^Mut^+ND: *P<0.05, **P<0.01, ***P<0.001, ns, not significant.

### Exacerbation of proteinuria and renal pathological changes in 60-week-old GCK^Mut^ mice fed with ND

Hyperglycemia is associated with microvascular complications in patients with T2D (17, 18). Given the elevated blood glucose in patients with GCK-MODY, it is important to determine the long-term effects of GCK-MODY on chronic diabetic complications. We therefore measured kidney weight (KW) and BW and found no significant difference in the KW/BW ratio between GCK^Mut^ and control mice at 28, 40, and 60 weeks of age (Fig. 2A). Additionally, in comparison to WT mice, GCK^Mut^ mice displayed no differences in the levels of 24-hour urinary albumin excretion (UAE), urinary albumin-to-creatinine ratio (UACR), and serum creatinine at 28 and 40 weeks of age. However, UAE, UACR, and serum creatinine were notably increased at 60 weeks of age in GCK^Mut^ mice (Fig. 2B-D). Histological analyses demonstrated relatively normal glomerular and tubular morphology and structure in GCK^Mut^ mice at 28 and 40 weeks of age. PAS and Masson staining indicated marginal differences in thylakoid dilation and renal fibrosis between GCK^Mut^ and control mice at 28 and 40 weeks, which exacerbated notably at 60 weeks of age (Fig. 2E-G and K-L). Subsequently, we utilized transmission electron microscopy to examine glomerular structure at 28, 40, and 60 weeks of age in both groups of mice. The observations indicated a trend of increased thickness of glomerular basement membrane (GBM) at 28 and 40 weeks of age, and a significant thickening of GBM was evident at 60 weeks of age in GCK^Mut^ mice (Fig. 2H-J and M).

**Figure 2.**
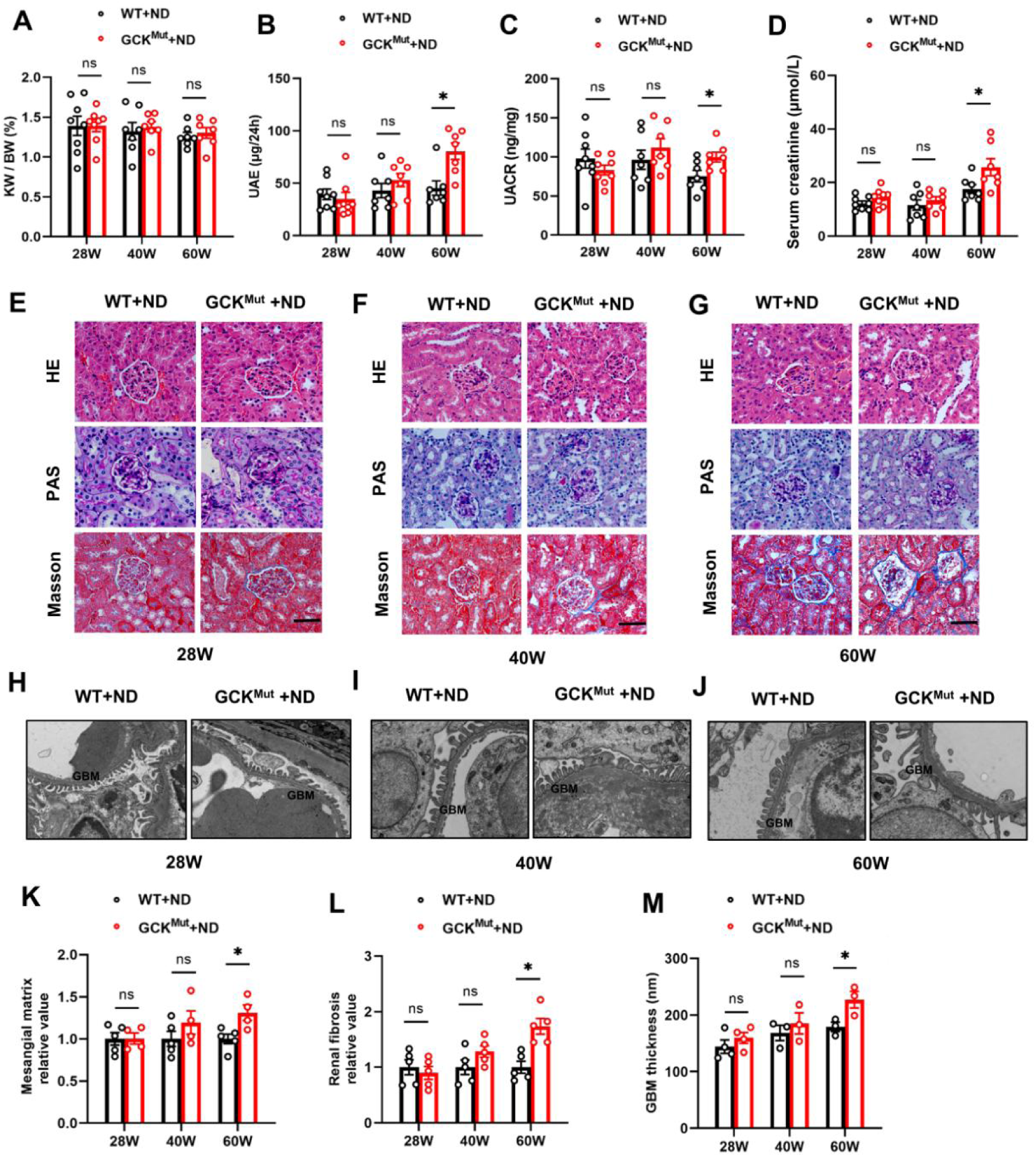
Changes in UAE, serum creatinine and renal histology in GCK^Mut^ mice fed with ND. (**A**) Kidney and body weights were measured at 28, 40, 60-week old, the ratio of kidney and bodyweight (KW/BW) were calculated. 24-hour UAE (**B**), UACR (**C**), and blood creatinine levels (**D**) were examined in two groups of mice fed with ND at 28, 40, and 60 weeks of age (n=7-8 mice/group). (**E-G**) H&E staining, PAS staining, and Masson staining of kidney sections were performed at 28, 40, and 60 weeks in mice fed with ND (n=5 mice/group). Original magnification = 400. Scale bar = 50μm. (**H-J**) Changes in glomerular structure observed by electron microscopy (n=3 mice/group). Original magnification is x8k. Semi-quantitative analysis of the mesangial matrix (**K**) and the degree of renal fibrosis (**L**). (**M**) Quantification of GBM thickness. Values are expressed as mean±SEM. WT+ND vs. GCK^Mut^+ND: *P<0.05, ns, not significant.

### Renal fibrosis- and inflammation-associated genes exhibit high expression in aging GCK^Mut^ mice fed with ND

To investigate pathways involved in long-term renal changes in GCK^Mut^ mice, we performed transcriptome analysis using kidneys from littermate controls and GCK^Mut^ mice. Gene Ontology (GO) analysis of the two groups of mice at 60-week-old age revealed that the 940 differentially expressed genes (DEGs, with │Log2│ fold change ≥1 and false discovery rate ≤0.05), which were primarily enriched in collagen fibril organization, cell-matrix adhesion, and inflammatory response (Fig. 3A), consistent with previous reports indicating that excessive accumulation of extracellular mesenchymal tissue and pro-inflammatory responses contribute to the progression of DKD (19–21). We further analyzed the DEGs involved in biological progression of DKD, and found that Vim (Vimentins, encoding intermediate filaments), Acta2 (α-SMA, encoding alpha-smooth muscle actin), Col1a1 (COL1A1, encoding type I collagen) and Tnf exhibited significant up-regulation in GCK^Mut^ mice at 60 weeks of age (Fig. 3B). The changes of the these genes were further confirmed by quantitative real-time PCR (qRT-PCR) excepted Colla1 that did not show a statistic difference between two groups. The gene Cdh1 (cadherin 1, encoding E-cadherin), which maintains stable cellular connections and cellular architecture integrity (22), was significantly downregulated (Fig. 3C). Furthermore, Western blot analysis verified that the protein levels of Vim, α-SMA, TNF-α and IL-1β were indeed significantly elevated in the kidneys of 60-week-old GCK^Mut^ mice, whereas no significant changes in proteins levels and mRNA levels were observed in 28- and 40-week-old mice (Fig. 3D-G, Fig. S2). These results suggest that upregulation of the genes associated with renal fibrotic and inflammatory responses may be attributed to long-term kidney injury in the old GCK^Mut^ mice.

**Figure 3.**
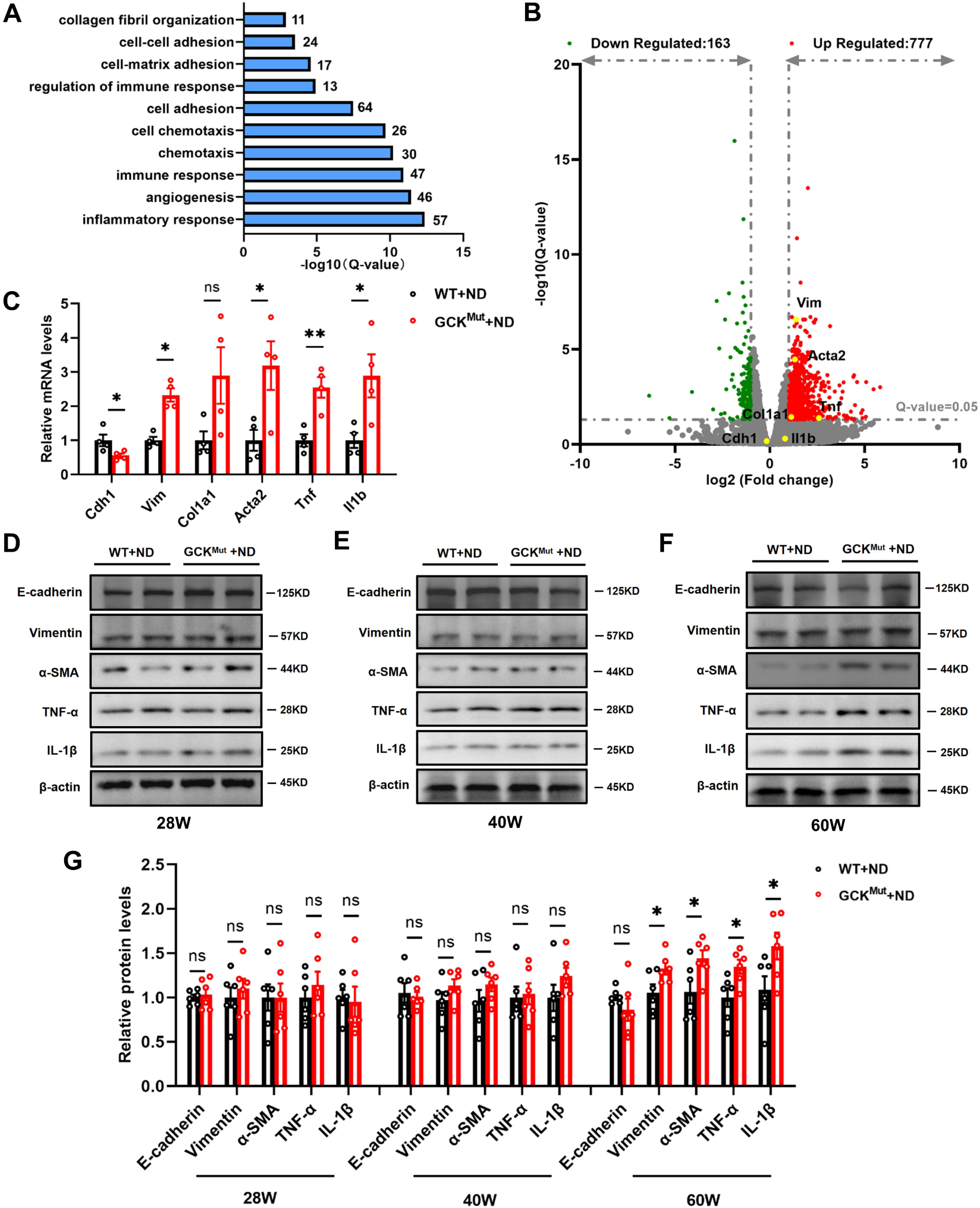
Increased renal fibrosis and inflammatory infiltration in 60-week-old GCK^Mut^ mice fed with ND. (**A**) GO analysis of RNA-seq data showed significant changes of pathways in the renal from 60-week-old WT control and GCK^Mut^ mice fed with ND (n=3mice/group). (**B**) Volcano plots depicting transcriptome data, dashed line on the y-axis indicates Q-value=0.05 and fold change on the x-axis is greater than 1. (**C**) The mRNA levels of indicated genes in the kidneys of 60-week-old WT control and GCK^Mut^ mice were determined by qRT-PCR (n=4 mice/group). (**D-F**) At 28, 40, and 60 weeks of age, the expression of E-Cadherin, Vimentin, α-SMA, TNF-α, and IL-1β in kidneys of control and GCK^Mut^ mice fed with ND were examined by western blots (n=6 mice/group). (**G**) Quantification of western blots shown in D-F. Values are expressed as mean±SEM. WT+ND vs. GCK^Mut^+ND: *P<0.05, **P<0.01, ns, not significant.

### The lipid profile of 28-week-old GCK^Mut^ mice fed with HFD improved, but worsened at 60 weeks

A recent study has revealed that partial inactivation of GCK can yield beneficial effects on hepatic lipid metabolism and inflammation (23). These effects have been postulated as a potential rationale behind the favorable lipid profile and reduced cardiovascular risk observed in patients with GCK-MODY (24, 25). Our data indicated that although improved circulating lipid profiles were indeed validated at early time in GCK^Mut^ Mutant mice, those improvements diminished over the time in mice fed with ND (Fig. 1O-R). We therefore asked what effects of additional metabolic stress inactivation of GCK on glucolipid metabolism and renal injury. We randomly grouped 4-week-old littermate Mutant and control mice for the HFD regimen while maintaining the same experimental protocols as those used for mice fed with ND. Our findings revealed a transient small but statistic decrease in body weight in 28-week-old GCK^Mut^ mice fed with HFD. However, this divergence disappeared in the mice fed with HFD till 40- and 60-week-old (Fig. 4A). Additionally, elevated fasting and postprandial blood glucose (Fig. 4B-E, Fig. S3A-C), decreased circulating insulin (Figs. 4F-H, Fig. S3D-F), and insulin sensitivity (Figs. 4I-K, Fig. S3G-I) had all similar trends as that we observed in mice fed with ND. These data further demonstrate that although elevated FBG and impaired glucose intolerance persist, these diabetes phenotypes are stable and are not aggravated even under HFD in GCK^Mut^ mice.

**Figure 4.**
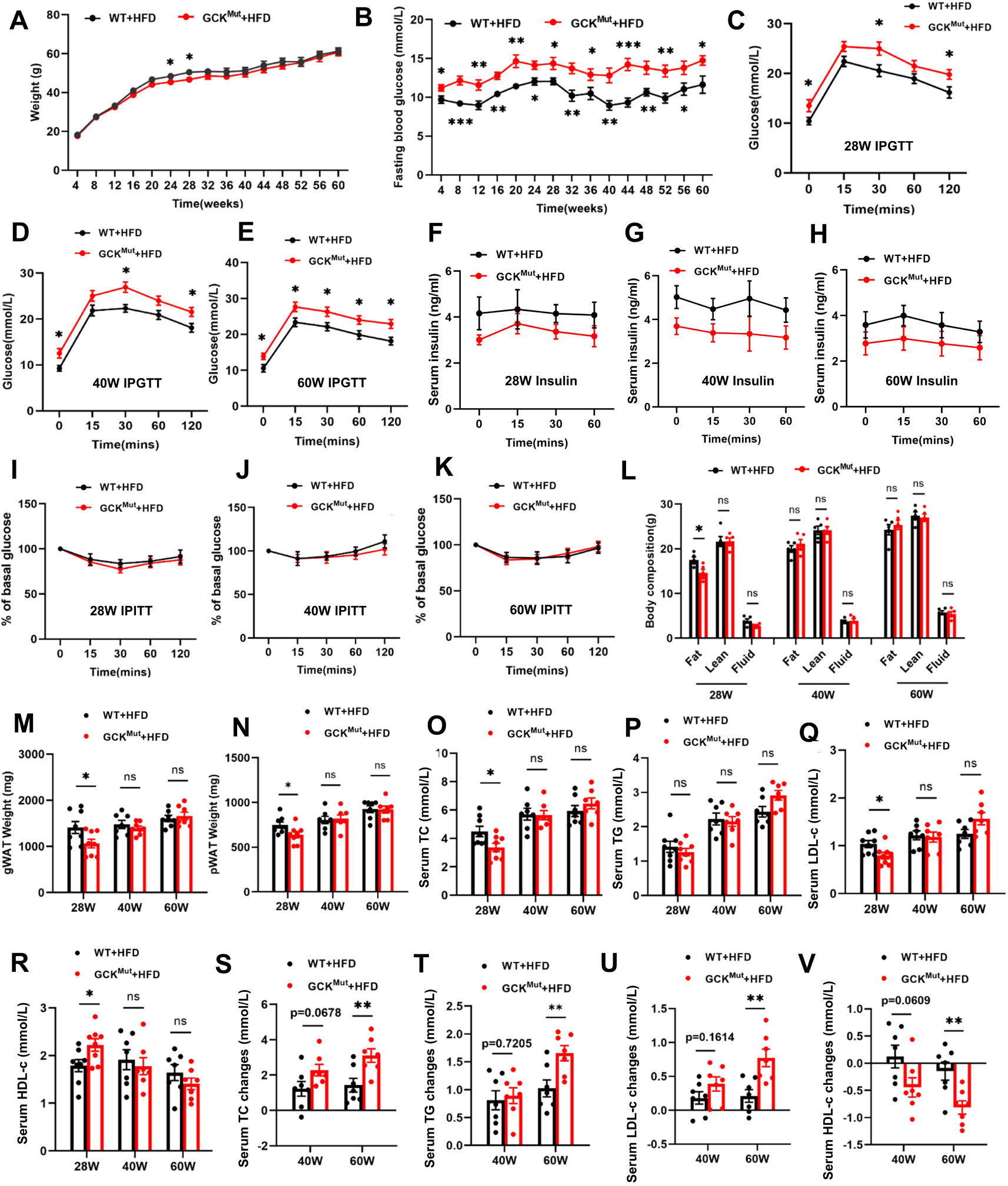
Changes in FBG and lipid metabolism in GCK^Mut^ mice fed with HFD. GCK^Mut^ and control mice were fed with HFD started from 4 weeks of age. The body weight (**A**) and FBG (**B**) have been monthly monitored and recorded till 60 weeks of age. IPGTT were performed in mice fed with HFD at 28 (**C**), 40 (**D**), and 60 (**E**) weeks of age. Serum insulin levels during IPGTT were also measured at 28 (**F**), 40 (**G**), and 60 (**H**) weeks of age. IPITT were performed in mice fed with HFD at 28 (**I**), 40 (**J**), and 60 (**K**) weeks of age (n=5-8 mice/group). (**L**) Body compositions were analyzed in the mice fed with HFD at 28, 40, and 60 weeks of age. (**M-N**) The weight of pWAT and gWAT were measured in the mice above. TC (**O**), TG (**P**), LDL-c (**Q**), HDL-c (**R**) were measured in the mice fed with HFD at 28, 40, and 60 weeks of age. (**S-V**) Changes of TC, TG, LDL-c, HDL-c in mice aged 40 weeks and 60 weeks compared with those aged 28 weeks. n=5-8 mice/group. Values were expressed as mean±SEM. WT+HFD vs. GCK^Mut^+HFD: *P<0.05, **P<0.01, ***P<0.001, ns, not significant.

Next, we analyzed body composition and revealed a significant reduction in fat mass in HFD fed GCK^Mut^ mice at 28 weeks in comparison to the control group (Fig. 4L). Similarly, pWAT and gWAT (Fig. 4M-N) as well as serum TC and LDL-c levels displayed noteworthy reductions along with elevated HDL-c in HFD fed GCK^Mut^ mice at 28 weeks (Fig. 4O-R). However, at 40 to 60 weeks of age, the aforementioned indices were progressively worse, resulting in significant deleterious changes of lipid profiles at the age of 60 weeks (Fig. 4S-V). These data suggest that long-term exposure of hyperglycemia and HFD may override the potential beneficial effects of GCK^Mut^ on lipid metabolism and body composition, manifesting more severe metabolism disorders.

### GCK^Mut^ attenuates HFD induced renal injury at 28 weeks of age, but aggravates at 60 weeks of age

Previous study has indicated that ectopic lipid accumulation in the kidneys contributes to kidney injury (26–28). Analogous to alterations in lipid profiles, quantitative analysis of lipids extracted from kidneys exhibited significantly reduced TC and TG levels along with decreased kidney weight in GCK^Mut^ mice fed with HFD at the age of 28 weeks. However, with extended exposure to the hyperglycemia and HFD, TC and TG contents in the kidneys of GCK^Mut^ mice gradually increased in comparison to WT mice over time (Fig. 5A-C, Fig. S4A-B). To further evaluate renal damage, we also examined 24-hour UAE, UACR, and serum creatinine levels in both groups of mice. The findings revealed reduced UAE, UACR, and serum creatinine levels in GCK^Mut^ mice fed with HFD at the age of 28 weeks, but increased alterations in 60-week-old HFD fed GCK^Mut^ mice (Fig. 5D-F, Fig. S4C-E). Notably, histological staining indicated significantly reduced renal mesangial matrix and collagen fiber deposition in GCK^Mut^ mice compared to WT mice fed with HFD at the age of 28 weeks, but these indicators gradually increased at 40 and 60 weeks (Fig. 5G-I and M-N). Furthermore, TEM analysis demonstrated a significantly smaller degree of GBM thickness in GCK^Mut^ mice at 28 weeks compared to control mice. However, this thickness significantly increased at 60 weeks (Fig. 5J-L and O). These findings indicate that proteinuria progression and pathological damage induced by a HFD were mitigated in GCK^Mut^ mice fed with HFD at early age, whereas long-term (60 weeks) HFD and hyperglycemia worsened renal damage at 60 weeks in GCK^Mut^ mice. Altogether, those findings highlight distinct diet and duration dependent effects of GCK inactivating mutation on lipid profiles and the kidney.

**Figure 5.**
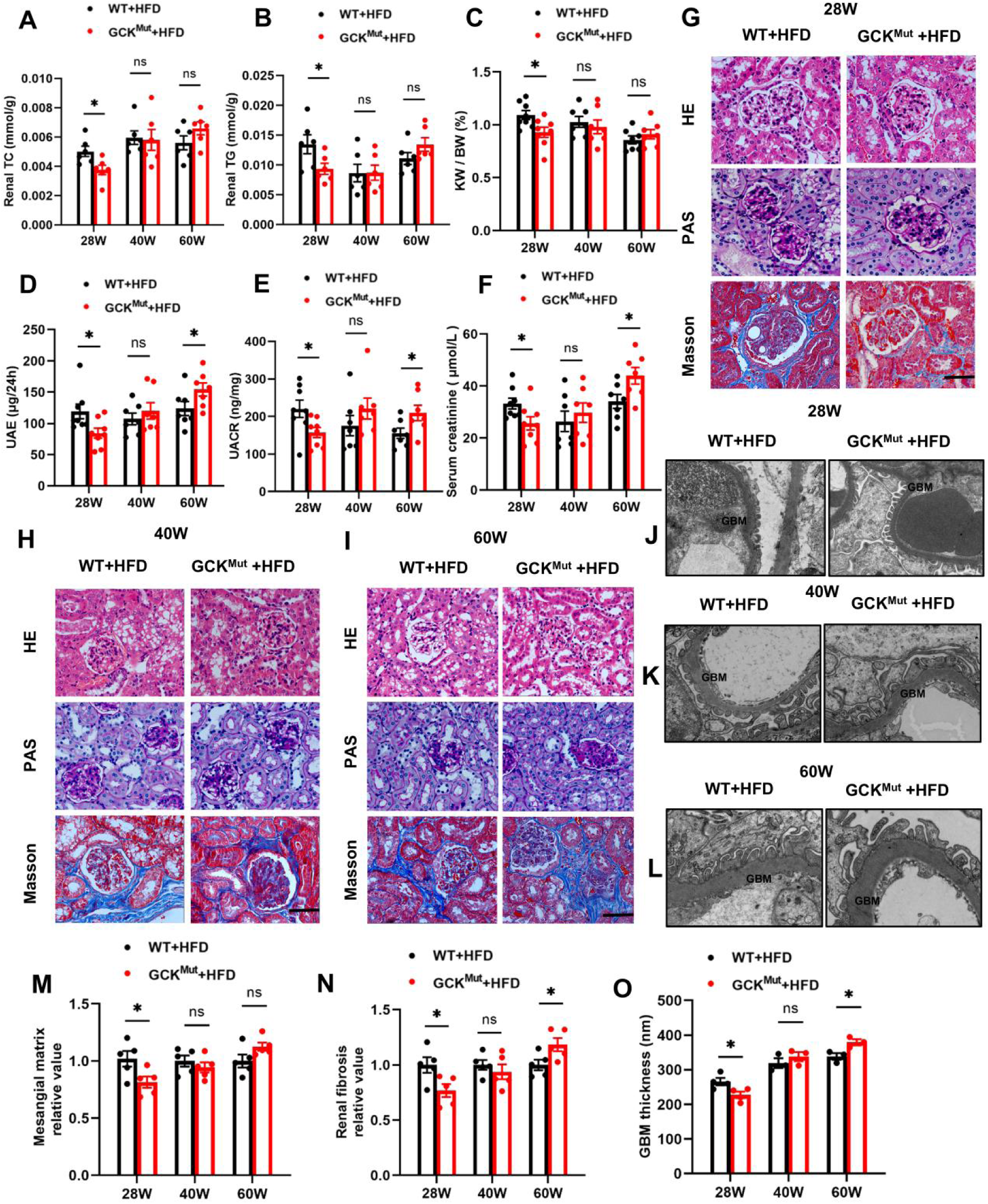
GCK^Mut^ attenuated HFD associated renal injury at 28 weeks of age, but aggravated renal injury at 60 weeks of age. Renal TC (**A**), Renal TG (**B**), KW/BW (**C**), 24-hour UAE (**D**), UACR (**E**), and serum creatinine levels (**F**) were examined at 28, 40, and 60 weeks of age in both groups of mice (n=7-8 mice/group). (**G-I**) H&E staining, PAS staining, and Masson staining of renal sections at 28, 40, and 60 weeks in both groups of mice (n=5 mice/group). Original magnification = 400. Scale bar = 50μm. (**J-L**) Electron microscopic observation of changes in glomerular structure in two groups of mice (n=3 mice/group). Original magnification is x8k. Semi-quantitative analysis of the mesangial matrix (**M**) and the degree of renal fibrosis (**N**). Quantification of GBM thickness (**O**). Values are expressed as mean±SEM. WT+HFD vs. GCK^Mut^+HFD: *P<0.05, ns, not significant.

### GCK^Mut^ attenuates renal fibrosis and inflammatory responses in 28-week-old mice fed with HFD, but exacerbates them at 60 weeks of age

To investigate the mechanism of attenuated renal injury in GCK^Mut^ induced by HFD, we conducted transcriptome analyses of renal samples from GCK^Mut^ mice fed with HFD at the age of 28 weeks. GO enrichment analysis revealed that 462 DEGs were predominantly involved in fatty acid metabolic process, extracellular matrix organization, immune system process, cell-matrix adhesion, collagen fibril organization and inflammatory response (Fig. 6A). Further analysis of genes enriched in GO highlighted that fibrosis-related genes such as Col1a1 and Vim, and inflammatory cytokines like Tnf were significantly down-regulated. Additionally, genes associated with lipid catabolic process, such as Ppara and Lpl, displayed significant up-regulation (Fig. 6B). Subsequently, qRT-PCR confirmed that mRNA levels of Vim, Tnf and Acta2 were significantly reduced and Cdh1, Ppara, whereas Lpl were significantly elevated in the kidneys of GCK^Mut^ mice compared with WT mice (Fig. 6C). Western blot also showed that the expression of E-Cadherin was increased, whereas the expression of Vim, α-SMA and TNF-α were decreased in GCK^Mut^ mice fed with HFD at the age of 28 weeks. Importantly, transcriptome analyses confirmed by qRT-PCR and western blotting showed that that the genes linked to renal fibrosis and inflammatory of the above gene expression were reversed in GCK^Mut^ mice fed with HFD at the age of 60 weeks (Fig. 6D-G, Fig. S5).

**Figure 6.**
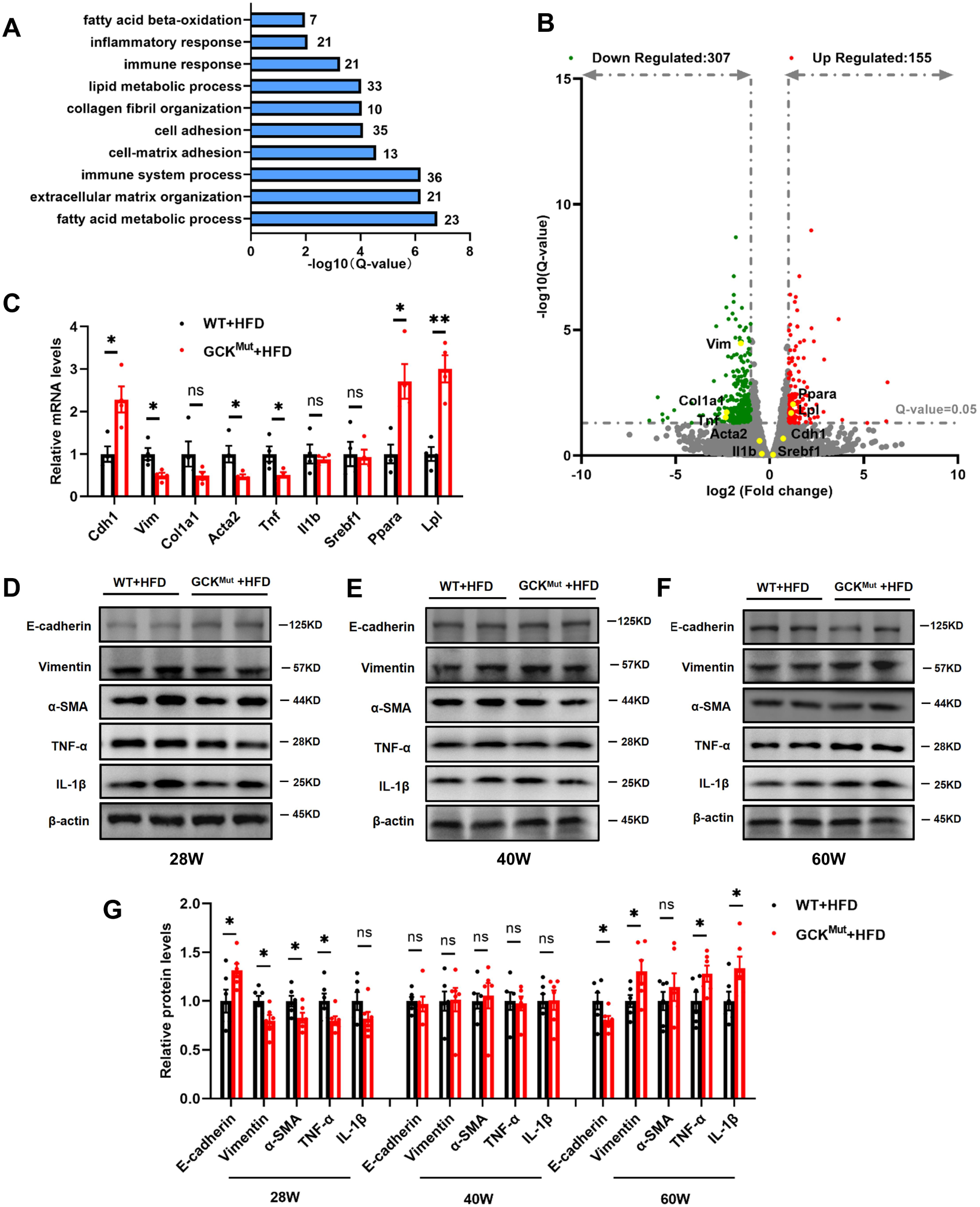
Renal fibrosis and inflammatory were attenuated in HFD GCK^Mut^ mice at 28 weeks of age and aggravated at 60 weeks of age. (**A**) The GO analysis of data obtained by RNA-Seq showed significantly different gene enrichment (n=3 mice/group). (**B**) Volcano plots depicting transcriptomics data, dashed line on the y-axis indicates Q-value=0.05 and fold change on the x-axis is greater than 1. (**C**) The mRNA levels of the mentioned factors in the kidneys of 28-week-old HFD WT control and GCK^Mut^ mice were determined by qRT-PCR (n=4 mice/group). (**D-F**) Renal E-Cadherin, Vimentin, α-SMA, TNF-α, and IL-1β protein levels were detected by Western blot analysis at 28, 40, and 60 weeks of age (n=6 mice/group). (**G**) Results of quantitative immunoblot analysis for each group. Values are expressed as mean±SEM. WT+HFD vs. GCK^Mut^+HFD: *P<0.05, **P<0.01, ***P<0.001, ns, not significant.

## Discussion

GCK-MODY is one of most common types of monogenic diabetes (29). Patients with GCK-MODY usually exhibit mild and asymptomatic fasting hyperglycemia, persisting from birth with minimal progression over time (4–6). Generally, the mild elevated blood glucose in GCK-MODY does not require treatment with antidiabetic medications, except during pregnancy (30–32). One primary rationale for this lies in the markedly low incidence of microvascular and macrovascular complications, nearly akin to those observed in healthy individuals (33, 34). Previous studies have revealed that patients with GCK-MODY showcase normal metabolic profiles, including relatively lower serum TG with higher HDL-c levels, which may contribute to fewer complications in these patients (35, 36). However, given the fact that uncontrolled prolonged hyperglycemia is strongly associated with diabetes complications and the lack of perspective studies focusing on patients with GCK-MODY in different dietary settings, the long-term effects of GCK inactivating mutations on chronic complications remain to be further determined.

In this study, using a newly established mouse model carrying a novel MODY causing GCK mutation, we delved into the impact of the GCK Mutant on lipid profiles and diabetic complications from youth to old age, paying particular attention to a commonly overlooked factors: the combination of mild hyperglycemia and a high-fat diet. We founded a reduction of abnormal lipid profiles and mitigated renal injuries in 28-week-old GCK^Mut^ mice fed with HFD. However, over an extended duration, a tendency towards exacerbated abnormalities in lipid metabolism and heightened renal damage was observed (Figs. 4-6). These data revealed surprisingly distinct duration and diet dependent effects of GCK^Mut^ and uncover previously unrecognized pathological significance of prolonged reduction of GCK activity in lipid metabolism and kidney injuries in patients with GCK-MODY. These data may be more informative as controlled experimental variables can minimize complex clinical factors including genetic diversity and use of different medications. These observations warrant further studies in humans.

One of clinical features of patients with GCK-MODY is optimal lipid profile characterizing as significantly lower levels of serum LDL-c and TG along with a higher level of HDL-c compared with nondiabetic control and patients with T2D (24, 36). A recent study reveals an increased recruitment of adipose triglyceride lipase and choline/ethanolamine phosphotransferase 1 onto the surface of HDLs, resulting in enhanced production of plasmalogen phosphatidylcholines that may confer vascular protection against chronic exposure to hyperglycemia (35). In this study, we found that although there were decreased levels of serum LDL-c and TG, there were no differences in serum HDL-c between control and GCK^Mut^ fed with ND (Fig. 1). However, under a metabolic stress (fed with HFD), GCK^Mut^ exhibits more beneficial effects not only on levels of HDL-c (Fig. 4), but also on renal lipid deposition and injuries (Fig. 5) at the age of 28 weeks. These observations suggest that beneficial effects GCK^Mut^ on lipid metabolism may override the negative impacts of hyperglycemia on the renal damage at the early stages of diabetes. However, as prolonged hyperglycemia further extends especially under the HFD conditions, these beneficial lipid metabolisms were gradually reversed along with increased proteinuria and renal injuries (Figs. 2-3 and 5-6). It remains to be determined whether these paradoxical duration and diet dependent effects of GCK^Mut^ were solely due to extra long-term exposure of hyperglycemia that directly disallows the protective effects of GCK^Mut^ on kidneys, or further prolonged hyperglycemia rejects beneficial effects of GCK^Mut^ on lipid metabolism, resulting more severe abnormal lipid profiles that may in turn damage kidney.

Recent studies have underscored the pivotal role of inflammation in the pathogenesis of DKD, indicating its significance in triggering metabolic, biochemical, and hemodynamic disturbances in diabetes (37, 38). DKD, recognized as a chronic, low-grade inflammatory disease, represents a primary feature associated with T2D and its microvascular complications (39–41). Renal inflammation manifests through immune cell infiltration, cytokine and chemokine release, and activation of inflammatory signaling pathways (42). Our investigation aligns with these findings, demonstrating that 60-week-old GCK^Mut^ mice, both on a ND or HFD, exhibited more severe renal inflammatory infiltration compared to controls (Figs. 3 and 6). Conversely, the observed reduction in renal proteinuria and pathological changes in 28-week-old GCK^Mut^ mice fed with HFD may be associated with diminished inflammatory infiltration. The characteristic pathological hallmark linked with DKD involves the accumulation of extracellular matrix (ECM) components in the glomeruli, often leading to renal angio-glomerulosclerosis and functional decline (19–22). Our study suggests that GCK^Mut^ may mitigate renal fibrosis by curbing ECM accumulation and expansion of the thylakoid matrix in the early stages of diabetic nephropathy, thereby substantially preserving renal function in diabetic mice. Additionally, previous research has indicated that kidneys of mice on a HFD exhibited notable glomerular and tubular injury, inclusive of substantial defects in the glomerular filtration barrier and heightened tubular cell apoptosis (43, 44). These effects were possibly linked to increased TC and TG content, alongside activation of the adipogenic pathway in the kidneys (45, 46). Correspondingly, our observations reinforce these findings, indicating that this protective effect on the kidney wanes or even exacerbates renal injury as the lipid profile deteriorates with prolonged high-fat feeding in GCK^Mut^ mice.

In summary, our study presents novel evidence emphasizing the pivotal role of GCK in glucolipid metabolism and the progression of diabetic complications. MODY causing GCK Mutant benefits lipid metabolism, decreasing renal lipid deposition, inflammatory infiltration, and fibrosis progression in the early stages, thus retarding disease advancement. However, GCK Mutant appeared to be associated with exacerbated abnormalities of lipid metabolism and renal injury in later stages. These distinct duration-dependent effects of GCK Mutant raise the question regarding current consensus for the management of patients with GCK-MODY.

## Materials and Methods

### Animal models

Male WT mice were purchased from SPF Biotechnology, while a global GCK-Q26L (GCK^Mut^) knock-in mouse line was established as described in our previous study. Both GCK^Mut^ mice and WT mice were housed in pathogen-free conditions within the the Laboratory Animal Center of Tianjin Medical University Chu Hsien-I Memorial Hospital. WT and GCK^Mut^ male mice were randomly assigned to four groups and subjected to either a normal diet (ND; comprising 20% protein, 4% fat, 5% fiber, SCXK(LU) 2018-0003) or a high-fat diet (HFD; comprising 20% protein, 60% fat, 20% carbohydrates, D12492, Research Diets) from four weeks of age until they reached 28, 40, and 60 weeks of age. Throughout this period, monitoring of body weight and blood glucose levels was conducted every 4 weeks. All mice utilized in the study were maintained on a C57BL/6J background and housed in a 12 h light/dark cycle in a temperature (22–25 °C) and humidity (55 ± 5%) control room.

### Biochemical measurements

Blood glucose levels were assessed using an automated glucometer (Roche, ACCU-CHEK, Germany). Mice were placed in metabolic cages to collect 24-hour urine samples. Various parameters including serum total cholesterol (TC), triglycerides (TG), low-density lipoprotein cholesterol (LDL-c), high-density lipoprotein cholesterol (HDL-c), serum creatinine, 24-hour urinary albumin excretion (UAE), and urinary creatinine were determined by kits provided from Nanjing Jiancheng Institute of Bioengineering (Nanjing, China).

### Intraperitoneal glucose tolerance test and insulin tolerance test

The intraperitoneal glucose tolerance test (IPGTT) involved the intraperitoneal administration of 1 g/kg body weight of glucose in mice following a 6-hour fasting period. The area under the curve (AUC) was determined using the trapezoidal rule. Simultaneously, blood samples collected at each time point were utilized to measure blood glucose and serum insulin levels employing an insulin ELISA assay kit (EZassay, China), following the manufacturer’s instructions. For the intraperitoneal insulin tolerance test (IPITT), mice that fasted for 4 hours received an intraperitoneal injection of 0.75 U/kg body weight of insulin. Tail vein glucose levels were monitored at 0, 15, 30, 60 and 120 minutes post-injection. All experiments mentioned above were conducted using age- and sex-matched cohorts.

### Body composition measurements

Fat mass of was measured using a Bruker Minispec mq7.5 (Brucker).

### Histologic evaluation and transmission electron microscopy anslysis

Following anesthesia, mice were euthanized subsequently the kidneys were harvested, which were fixed in 4% paraformaldehyde, then paraffin-embedded and were cut into 5-μm-thick sections. Subsequently, these sections were stained, and after sealing with neutral balsam, images were captured using AxioImagerM2 (Carl Zeiss, Oberkochen, Germany). The extent of kidney injury within each group was assessed utilizing Image-ProPlus 6.0 software. The sections underwent staining procedures involving hematoxylin and eosin (H&E), Periodic acid-Schiff’s (PAS), and Masson’s trichrome. For semi-quantitative analysis, twenty stained glomeruli per section were randomly chosen (5 mice per group) for analyses. For transmission electron microscopy (TEM), fresh isolated kidney tissues were sliced into small pieces and then fixed in 2.5% glutaraldehyde for a duration of 12 hours. Subsequently, the tissues underwent treatment with 1% osmium tetroxide. Ultrathin sections were obtained utilizing an ultramicrotome (LeicaEMUC7, Germany), followed by a dehydration process using gradient alcohol, permeabilization with acetone, and final embedding in epoxy. The obtained ultrathin sections, measuring between 70-90 nm, were stained with uranyl acetate and lead citrate. These sections were examined using a transmission electron microscope (TEM) (HT7700, Japan).

### RNA Extraction, cDNA Synthesis, and Quantitative real-time PCR (qRT-PCR)

Total RNA was extracted and purified from renal tissues using TRIzol reagent (Invitrogen, USA). Subsequently, cDNA was synthesized utilizing the PrimeScriptTM RT kit with gDNAEraser (Takara, Japan), following the manufacturer’s instructions. qRT-PCR assays were conducted in a 20 μL reaction volume employing TB Green Premix Ex Taq^TM^ (Tli RNaseH Plus) (Takara, Japan). GAPDH served as the internal control. The primers utilized in this study were designed and chemically synthesized by ZTSINGKE Biological Technology (Beijing, China) Co. Ltd. Details of the primer sequences were provided in **Supplemental Table S1**.

### Protein Extraction and Western Blot Analysis

Renal tissues were lysed using RIPA lysis buffer for protein extraction. Protein samples were then loaded onto SDS-PAGE, electrophoretically separated, and subsequently transferred to nitrocellulose membranes. These membranes were blocked using 5% skimmed milk followed by blotted with indicated antibodies, and visualized by enhanced chemiluminescence (CLiNX, Shanghai, China). The following antibodies were utilized in this study: anti-E-Cadherin (A3044; ABclonal), anti-Vimentin (A11423; ABclonal), anti-α-SMA (A7248; ABclonal), anti-IL-1β (A16288; ABclonal), anti-TNF-α (A11534; ABclonal), and anti-β-actin (KM9001T, Sungene).

### Transcriptome analysis

Renal transcriptome analysis was conducted by BGI Group, encompassing RNA extraction, library preparation, and sequencing. Data processing adhered to the standard BGI mRNA analysis pipeline. Statistical analysis was executed utilizing the Deseq2 software package (v1.28.1). Differentially expressed genes (DEGs) were identified using a threshold of log2 (fold change) absolute value ≥ 1 and false discovery rate ≤ 0.05. For further insights, Gene Ontology (GO) analysis was carried out using cluster/profiler (v3.16.1). Pathway visualization was accomplished using GraphPad Prism 8.

### Statistics

Statistical analyses were conducted utilizing GraphPad Prism 8 software. The data are presented as mean ± standard deviation. Analysis involved unpaired Student’s t-tests, with statistical significance set at the traditional critical value of P < 0.05. Each experiment was independently repeated at least three times.

## Data availability

The authors are willing to make the raw data generated in their study available to other investigators upon reasonable request.

## Author contributions

YH, YF, YF, XL, XL,WQ and YY conducted experiments and analyzed data; YL, WL, ZZ, SM, SJ, SC, HS, KW and WF conducted experiments; QH discussed project and edited manuscript; ML initiated and oversaw the project, designed the experiments, wrote the manuscript, and is responsible for the integrity of the data and the accuracy of the data analysis.

## Acknowledgments

The work was supported by the National Key R&D Program (2022YFE0131400 and 2019YFA0802502), and also by the National Natural Science Foundation of China (82220108014, 82100865, and 82301951). We acknowledge the support of Tianjin Key Medical Discipline (Specialty) Construction Project (TJYXZDXK-030A), Tianjin Municipal Health Commission (TJWJ2021ZD001), Tianjin Municipal Science and Technology Commission (23JCQNJC00680), Tianjin Medical University General Hospital Clinical Research Program (22ZYYLCZD02), and Tianjin Medical University Clinical Special Disease Research Center - Neuroendocrine Tumor Clinical Special Disease Research Center.

---

**Supplemental Figure 1.**
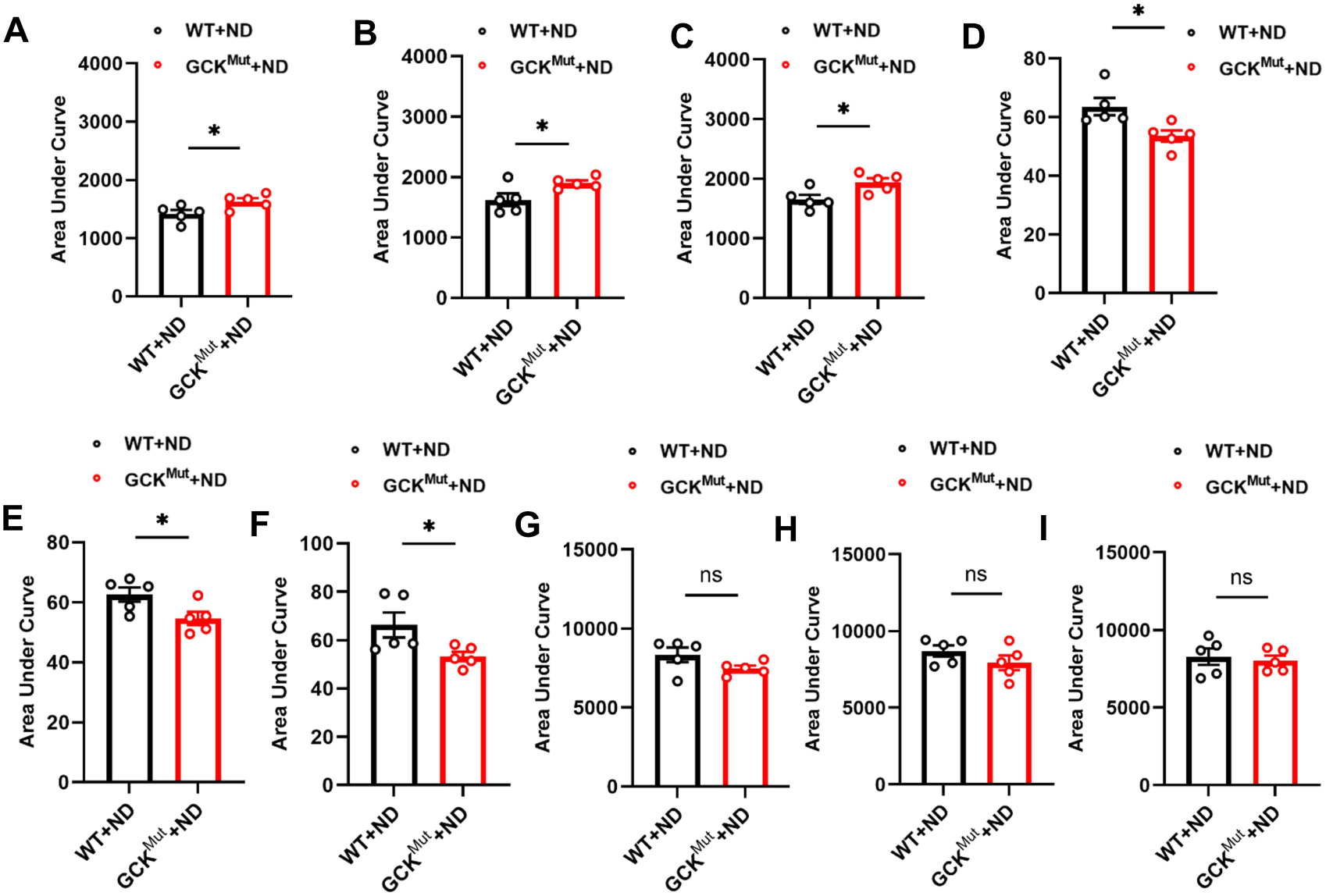
The area under the curve (AUC) of IPGTT, serum insulin and IPITT in two groups of mice on ND. (**A-C**) The AUCs of blood glucose levels were calculated according to Fig.1C-E. (**D-F**) The AUCs of serum insulin levels were calculated from Fig. 1F-H. (**G-I**) The AUCs of blood glucose during IPITT were calculated from Fig.1I-K. n=5mice/group. Values are expressed as mean±SEM. WT+ND vs. GCK^Mut^+ND: *P<0.05, ns, not significant.

**Supplemental Figure 2.**
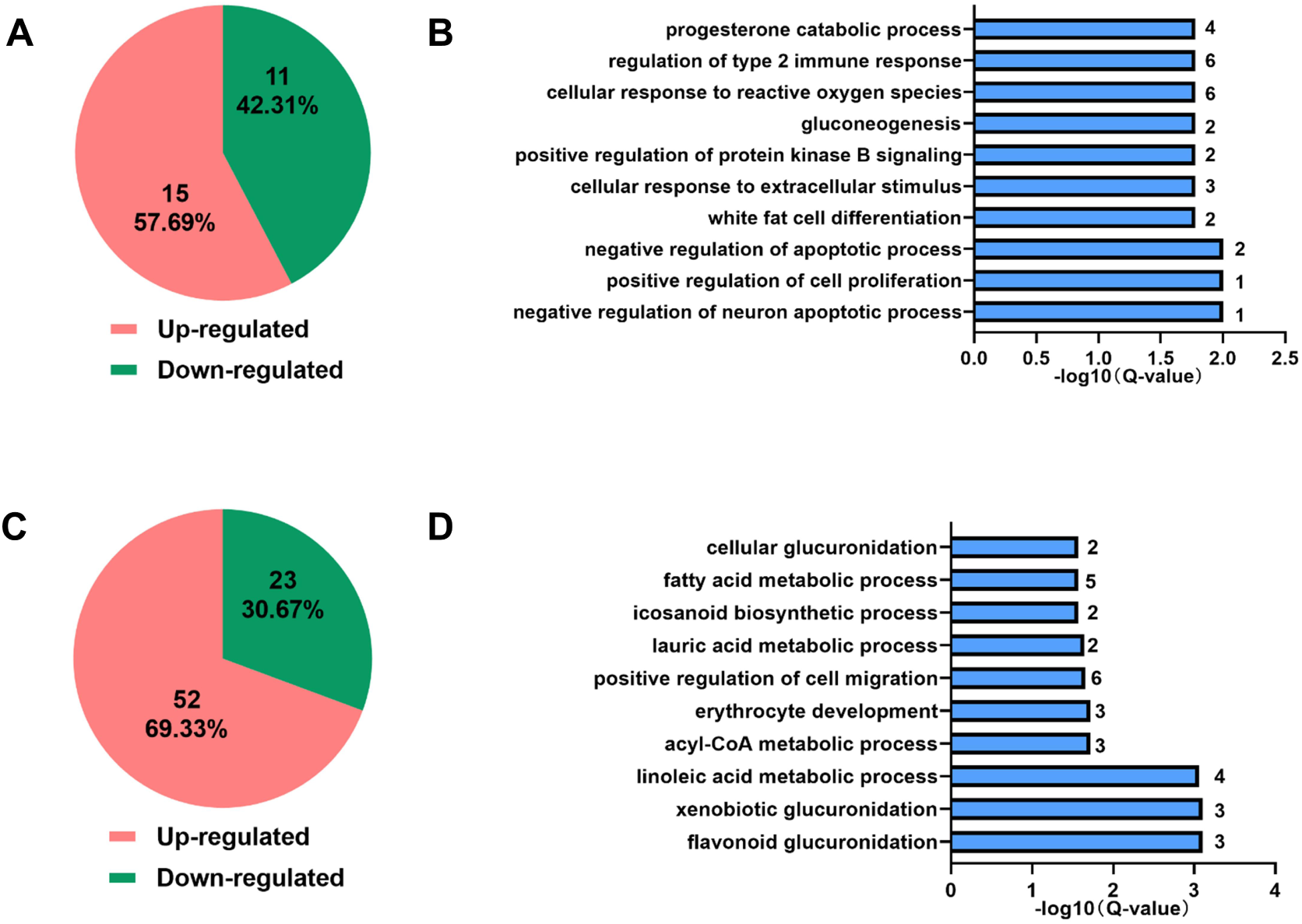
The kidney transcriptome analysis at 28 and 40 weeks in two groups of mice on ND. (**A-B**) The number of significantly different genes up- and down-regulated and GO analysis in two groups of mice on regular diet at 28 weeks of age. (**C-D**) The number of significantly different genes up- and down-regulated and GO analysis in two groups of mice on ND at 40 weeks of age. n=3 mice/group.

**Supplemental Figure 3.**
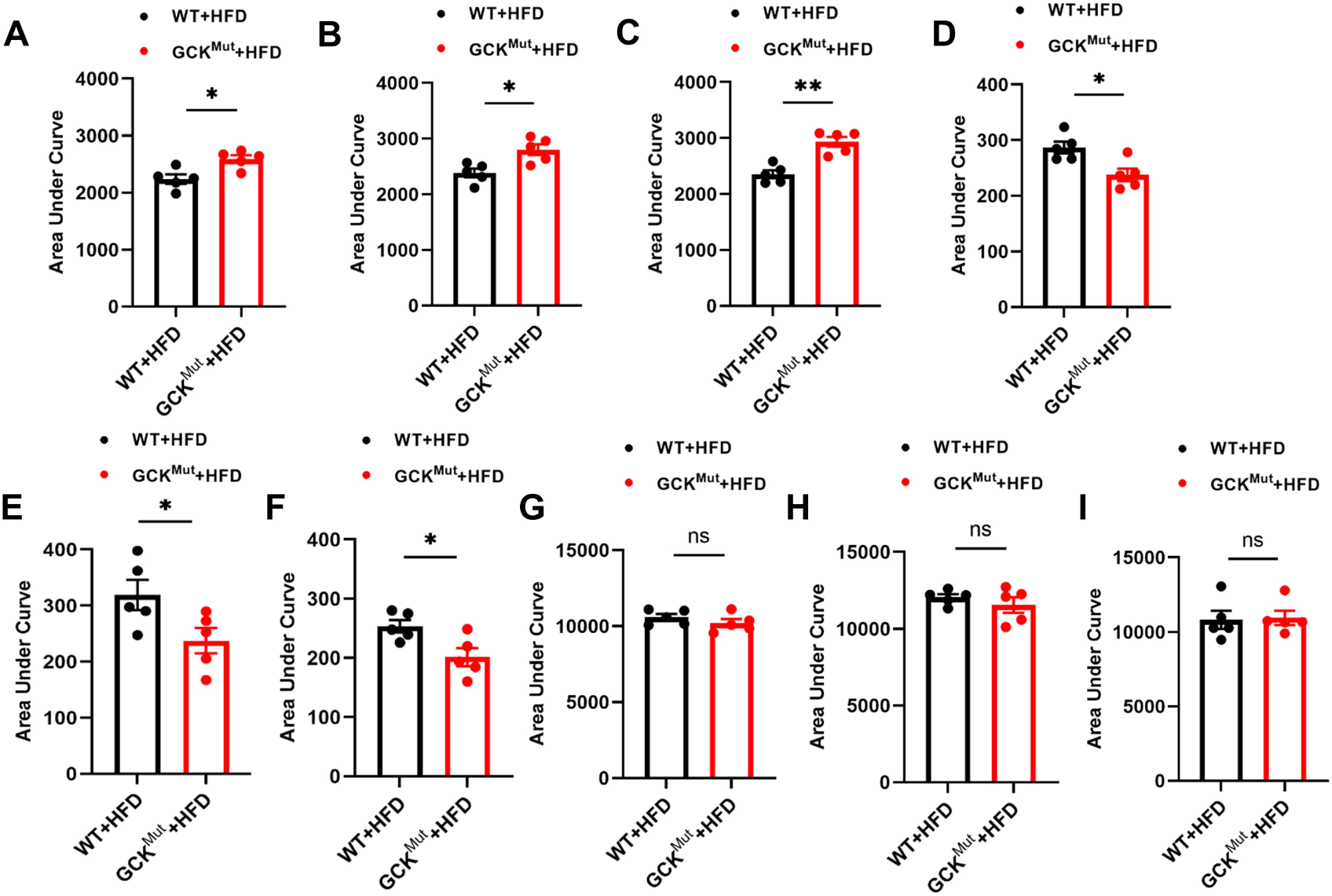
The area under the curve (AUC) of IPGTT, serum insulin, and IPITT in two groups of mice fed with HFD. (**A-C**) The AUC of IPGTT was calculated according to Figure.5C-E. (**D-F**) The AUC for serum insulin levels was calculated from Figure.5F-H. (**G-I**) The AUC values for IPITT were calculated from Figure.5I-K. n=5 mice/group. Values are expressed as mean±SEM. WT+HFD vs. GCK^Mut^+HFD: *P<0.05, ns, not significant.

**Supplemental Figure 4.**
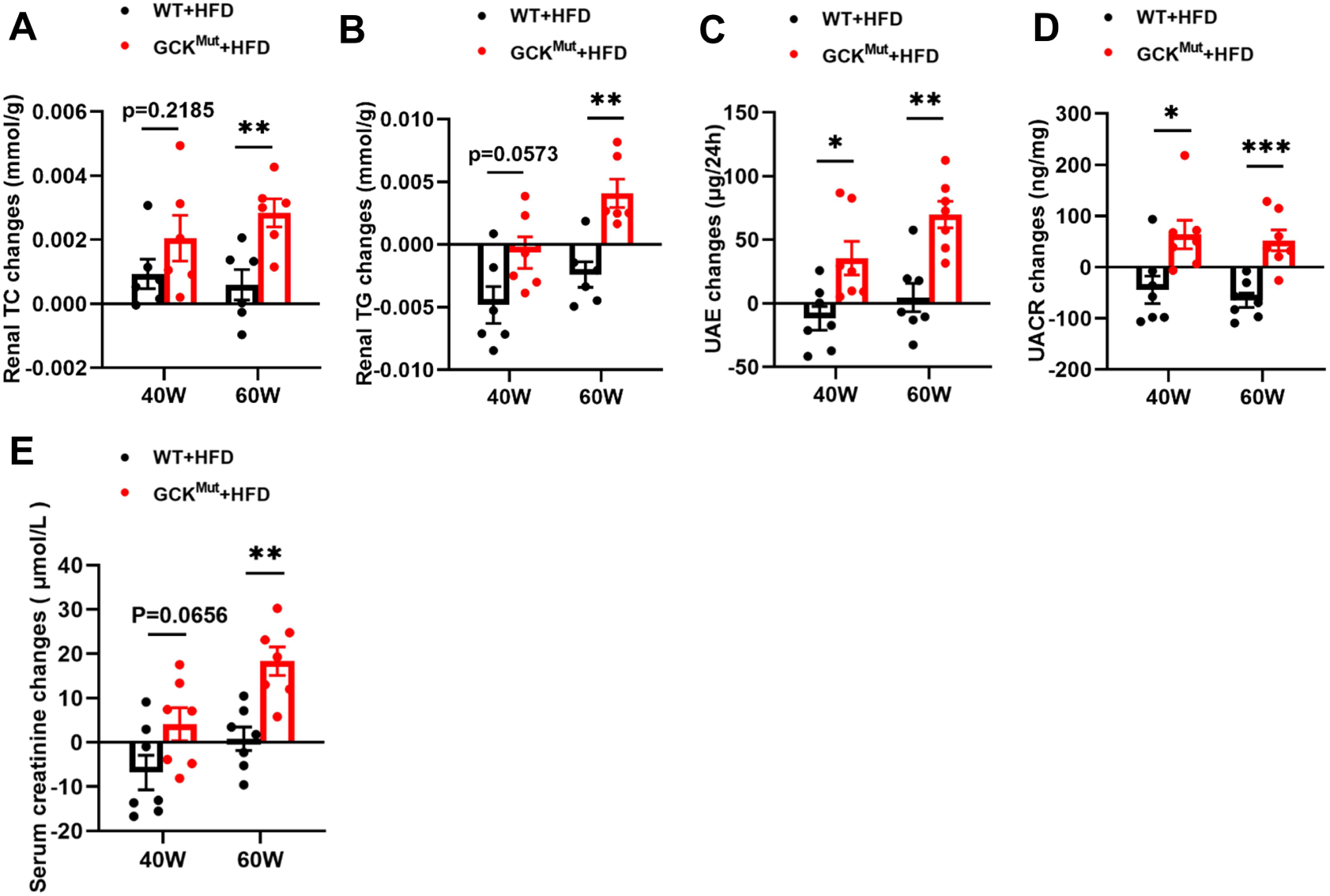
Changes of Renal TC, Renal TG, 24-hour UAE, UACR, and serum creatinine in mice aged 40 weeks and 60 weeks compared with those aged 28 weeks fed with HFD. Changes of Renal TC (**A**), Renal TG (**B**), 24-hour UAE (**C**), UACR (**D**), and serum creatinine (**E**) in mice aged 40 weeks and 60 weeks compared with those aged 28 weeks. Values were expressed as mean±SEM. WT+HFD vs. GCK^Mut^+HFD: *P<0.05, **P<0.01, ***P<0.001, ns, not significant.

**Supplemental Figure 5.**
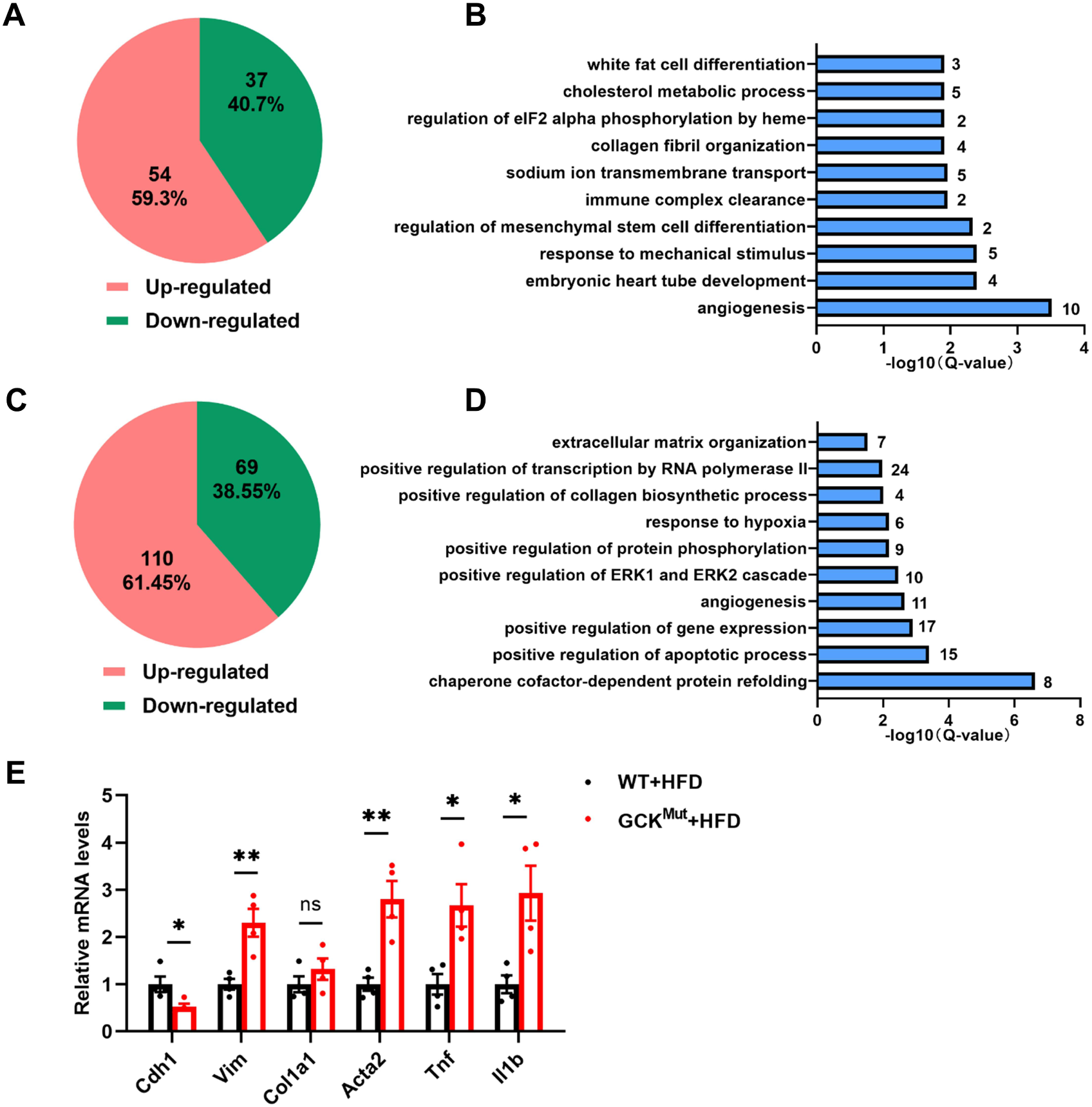
The kidney transcriptome analysis at 40 and 60 weeks in two groups of mice on HFD. (**A-B**) The number of significantly different genes up- and down-regulated and GO analysis in two groups of mice on HFD at 40 weeks of age. (**C-D**) The number of significantly different genes up- and down-regulated and GO analysis in two groups of mice on HFD at 60 weeks of age. (**E**) The mRNA levels of relevant fibrosis and inflammatory factors in the kidneys of two groups of mice on HFD at 60 weeks of age.

**Supplemental Table 1.**
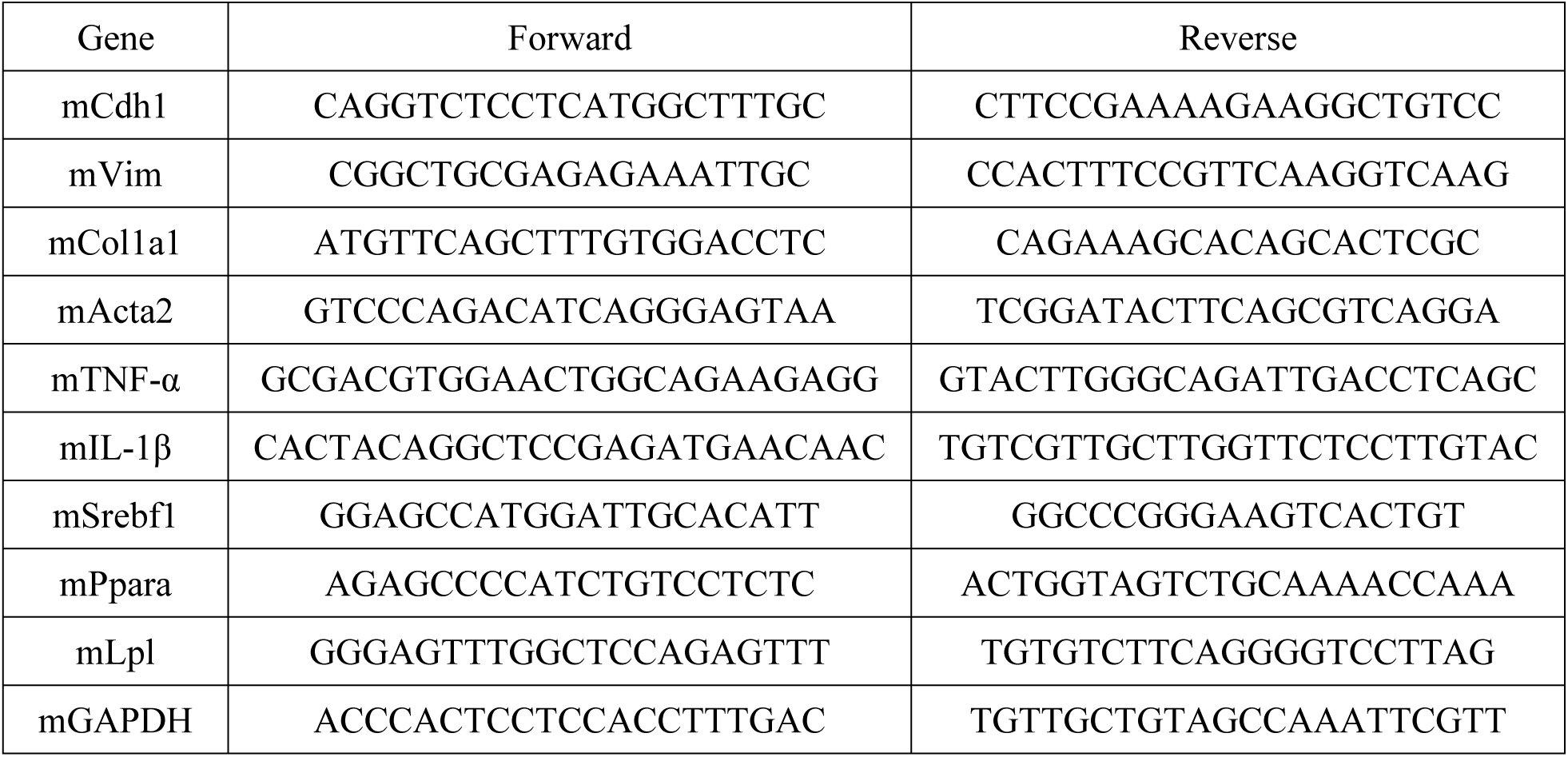
Primer sequences (m-Mus musculus) for qRT-PCR (5’-3’)

